# Ultra-sensitive sequencing for cancer detection reveals progressive clonal selection in normal tissue over a century of human lifespan

**DOI:** 10.1101/457291

**Authors:** Jesse J. Salk, Kaitlyn Loubet-Senear, Elisabeth Maritschnegg, Charles C. Valentine, Lindsey N. Williams, Reinhard Horvat, Adriaan Vanderstichele, Daniela Nachmanson, Kathryn T. Baker, Mary J. Emond, Emily Loter, Thierry Soussi, Lawrence A. Loeb, Robert Zeillinger, Paul Speiser, Rosa Ana Risques

## Abstract

High accuracy next-generation DNA sequencing promises a paradigm shift in early cancer detection by enabling the identification of mutant cancer molecules in minimally-invasive body fluid samples. We demonstrate 80% sensitivity for ovarian cancer detection using ultra-accurate Duplex Sequencing to identify *TP53* mutations in uterine lavage. However, in addition to tumor DNA, we also detect low frequency *TP53* mutations in nearly all lavages from women with and without cancer. These mutations increase with age and share the selection traits of clonal *TP53* mutations commonly found in human tumors. We show that low frequency *TP53* mutations exist in multiple healthy tissues, from newborn to centenarian, and progressively increase in abundance and pathogenicity with older age across tissue types. Our results illustrate that subclonal cancer evolutionary processes are a ubiquitous part of normal human aging and great care must be taken to distinguish tumor-derived, from age-associated mutations in high sensitivity clinical cancer diagnostics.

## INTRODUCTION

Worldwide more than a quarter of a million new cases of ovarian cancer are diagnosed each year and two thirds of these women die from the disease (*1*). This high mortality is largely due to the high frequency of metastasis before symptoms lead to diagnosis and a lack of effective screening methods. More than 60% of cases are diagnosed at an advanced stage, when the 5-year survival rate is only 29% (*2*). In contrast, survival for women with localized disease is 92%, indicating that early ovarian cancer detection could vastly decrease mortality, yet diagnosis at this stage is rare.

Despite significant efforts, unlike colonoscopies, pap smears and mammograms that are evidence-based, mortality-reducing methods for early detection of colon, cervical and breast cancer, respectively, no ovarian cancer screening technique has demonstrated sufficient sensitivity and specificity for use in the general population (*3*). The approach most explored involves a combination of testing for the serum protein CA-125 and transvaginal ultrasound, but the US Preventive Services Task Force recommends against its use (*4*) because it has not been shown to reduce mortality and may result in harms due to false positives, such as unnecessary surgeries in cancer-free women (*5*). Better tools for early ovarian cancer detection remains an urgent and unmet clinical need.

In the last several years an increasing number of cancers have been found to shed cells or DNA into blood or other body fluids where they can be non-invasively detected, a concept often termed “liquid biopsy” (*6*). Proof-of-principle for this approach in ovarian cancer screening was initially accomplished via identification of tumor-derived mutations in DNA extracted from routine Pap smear fluid (*7*). Although the sensitivity for ovarian cancer detection was only 41%, these findings supported the exciting possibility that ovarian cancer could be detected based on the genetic identification of cancer cells disseminated into the gynecological tract. A follow-up study recently reported that improved sensitivity, up to 63%, could be obtained by combining mutation detection in Pap tests and plasma. In addition, sampling with an intrauterine brush also improved sensitivity, probably due to increased tumor cell recovery by more proximal collection to the anatomical site of tumors (*8*).

An alternate means for tumor cell collection, developed by members of our team several years ago, consists on trans-cervical lavage of the uterine cavity (**Fig. 1A**). This method improves the efficiency of collection by rinsing all surfaces, including those near, and even into, the fallopian tubes where most early serous ovarian cancers are believed to initiate (*9*). This lavage technique demonstrated 80% sensitivity for ovarian cancer detection. The challenge, however, was that cancer-derived mutations, particularly those from early stage tumors, were often present in a very small fraction of the total lavage DNA. To detect these mutations, digital droplet PCR (ddPCR) was required, which is an extremely sensitive method but not practical for prospective screening because the tumor mutation needs to be known *a priori*.

**Figure 1.**
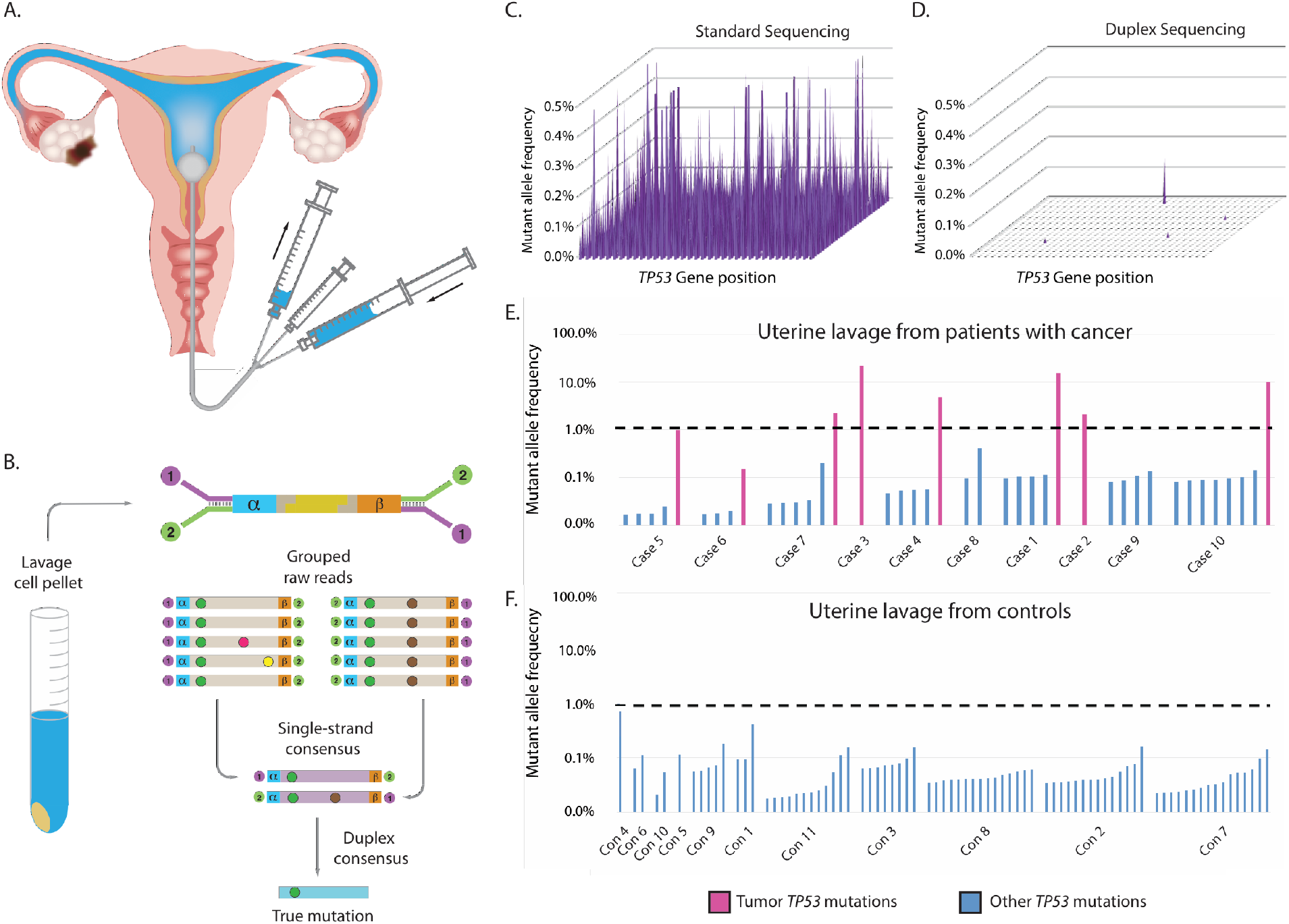
Detection of ovarian cancer using uterine lavage plus Duplex Sequencing. (A) Uterine lavage is carried out by passing a small catheter through the cervix followed by concurrent flushing and aspiration with 10 mL of saline as described (*24*). (B) After cell isolation by lavage centrifugation, DNA is extracted, fragmented, and ligated with specialized Duplex Sequencing adapters that include degenerate molecular tags (*α* and *β*). Following amplification, hybrid capture and sequencing, reads sharing the same barcodes are grouped into families and mutations are scored only if present in both strands of each original DNA molecule. (C) Each spot on the 2 dimensional surface represents one of the 1179 coding positions in *TP53*. The height of each peak indicates the mutant allele frequency (MAF) at each position as determined by conventional NGS, which shows false mutations at every position. (D) DS of the same sample (case 6 below) eliminates errors and reveals only true mutations. (E) *TP53* mutations identified by DS in uterine lavage from women with ovarian cancer and (F) cancer-free (controls). Fuchsia bars represent the matching tumor mutation and blue bars represent ‘biological background’ mutations. Mutations are sorted by ascending MAF within each patient and patients are sorted by age. Dashed lines indicate the optimal threshold to distinguish patients with and without ovarian cancer (sensitivity: 70%, specificity: 100%).

Next-generation DNA sequencing (NGS) is a widely used, variant-agnostic form of mutation detection, but has a background error rate of up to ~1%, which precludes confident detection of lower frequency mutations (*10*). Of the mutations comprising the 80% sensitivity achieved in our previous study, conventional NGS missed approximately one quarter (*9*). Currently, the most sensitive NGS method is Duplex Sequencing (DS) (*10*), which employs double-stranded molecular barcodes for error correction and decreases the error rate of sequencing from 10^-3^ to <10^-7^ (*11, 12*).

We previously demonstrated that DS can detect ovarian cancer-derived mutations in DNA extracted from peritoneal fluid at frequencies as low as one tumor mutation per 25,000 normal genomes (*13*). The extreme sensitivity of the technique also led to the discovery of prevalent, yet very low frequency (<0.01%) *TP53* mutations in both the peritoneal fluid and peripheral blood from healthy women. These ‘biological background’ mutations resembled *TP53* mutations found in cancers, but appeared to result from the normal aging process. This observation was among the first of an emerging body of literature that has identified age-related, cancer-associated mutations within non-cancerous tissue (*14*).

In the present study, we combine the most sensitive reported sampling technique for ovarian cancer detection, uterine lavage, with the highest accuracy sequencing technology available, DS. High-grade serous ovarian carcinoma (HGSOC) is both the most common and most deadly histological type, accounting for 70-80% of ovarian cancer deaths (*15*). With this in mind, we focus our sequencing efforts solely on *TP53* because >98% of HGSOCs carry an inactivating mutation in this relatively small and well-characterized gene (*16, 17*) and because *TP53* mutations have been routinely found in microscopic carcinoma-*in-situ* precursor lesions in the fallopian tubes, indicating that they are among the earliest genetic events in HGSOC formation (*18, 19*).

The primary goal of this study is to demonstrate the clinical and technical feasibility of using DS to deeply sequence *TP53* from uterine lavage as a potential screening test for ovarian cancer detection. We capitalize on the extreme sensitivity of ultra-accurate DS to identify cancer-derived mutations as well as to uniquely detect low frequency, age-related mutations that might impact diagnostic specificity. To better understand the extent and nature of these ‘biological background’ mutations, we perform a detailed characterization of somatic *TP53* mutations in multiple gynecologic tissues from women without ovarian cancer of ages spanning a century of human lifetime.

## RESULTS

### Study design and technology rational

Prior work by members of our group demonstrated 80% sensitivity to detect ovarian cancer, including small volume, early stage disease, by using a combination of uterine lavage and mutation analysis via massively parallel sequencing and ddPCR (*9*). While highly sensitive, ddPCR requires baseline knowledge of the specific mutations sought and thus is not practical for cancer screening. In the current study, we tested whether ultra-high accuracy DS could identify ovarian cancer mutations in uterine lavage with sensitivity comparable to ddPCR, but without prior knowledge of the tumor mutation.

We used DS to examine the coding region of *TP53* in DNA extracted from the lavage cell pellet of 10 women with ovarian cancer and 11 controls under blinded conditions. Most of these samples were included in the original study (*9*) (**Table S1**). DS employs special adapters with double-stranded molecular barcodes, which allow the identification of sequencing reads that were derived from both strands of each starting DNA molecule. Mutations are only scored if they are present in the majority of reads from both DNA strands, effectively eliminating sequencing and PCR artifacts (**Fig. 1B**). The estimated error rate of DS is below one-in-ten-million (*11*), which allows for extreme sensitivity and specificity of mutation detection when carrying out high depth sequencing (**Fig. S1**).

### Duplex Sequencing detects ovarian cancer mutations in uterine lavages with high sensitivity

To illustrate the superior accuracy of DS compared with standard Illumina sequencing, an example of a uterine lavage sample (case 6) processed by both methods is shown side-by-side in **Fig. 1C-D**. Whereas every nucleotide position in the gene artefactually appears mutated in 0.1-1% of molecules with standard sequencing, DS eliminates these tens of thousands of erroneous mutations to reveal the known tumor mutation at a mutant allele frequency (MAF) of 0.15%, a value very close to the frequency previously determined by ddPCR (0.12%, case 6 in **Table 1**) (*9*).

Among the 10 lavages from women with ovarian cancer, we identified the expected tumor mutation (fuchsia bars) in 8, matching the 80% sensitivity of the previous study (**Fig. 1E**). In the subset of these lavages that had been analyzed by conventional NGS and/or ddPCR, we confirmed tumor mutations at similar allele frequencies in most cases (**Table 1**, top). In addition to the tumor mutations, in nearly all lavages from women with and without tumors we identified very low frequency *TP53* mutations (blue bars) (**Fig. 1F**). To confirm that these mutations were not due to technical errors, two of the mutations identified in controls (lavages con2 and con7) were assessed by ddPCR (**Table 1**, bottom). This orthogonal assay demonstrated that these mutations, present at a comparable frequency <0.1% by both assays, were authentic.

**Table 1.** Comparison of *TP53* mutant allele frequencies by standard NGS, Duplex Sequencing, and digital droplet PCR

| Patient ID | Age | Histology | FIGO | Tumor mutation-DNA | Tumor mutation-protein | Mutant Allele Frequency of tumor *TP53* mutations |  |  |
| --- | --- | --- | --- | --- | --- | --- | --- | --- |
|  |  |  |  |  |  | Standard NGS MAF | Duplex Sequencing MAF | ddPCR MAF |
| case 1 | 67 | HGSOC | IIIC | c.578A>C | p.H193P | 15.25% | 15.63% | 14.45% |
| case 2 | 69 | HGSOC | IIIB | c.646G>A | p.V216M | 2.22% | 2.12% | n.a. |
| case 3 | 51 | Signet ring<sup>†</sup> | na | c.844C>T | p.R282W | 22.76% | 22.22% | 25.55% |
| case 4 | 54 | HGSOC | IV | c.785G>T | p.G262V | 5.05% | 4.88% | n.a. |
| case 5 | 32 | HGSOC | IIIC | c.743G>A | p.R248Q | n.a. | 0.98% | 0.07% |
| case 6 | 42 | HGSOC | IIIA | c.503A>G | p.H168R | Negative | 0.15% | 0.12% |
| case 7 | 55 | HGSOC | IIIA | c.815A>T | p.V272E | Negative | Negative | n.a. |
| case 8 | 48 | HGSOC | IIIC | c.734G>T | p.G245V | Negative | 2.26% | 0.20% |
| case 9 | 67 | HGSOC | IV | c.1024C>T | p.R342* | Negative | Negative | n.a. |
| case 10 | 68 | HGSOC | IIIC | c.535C>T | p.H179Y | n.a. | 9.95% | n.a. |
| Patient ID | Age | Histology | FIGO | Non cancer mutation-DNA | Non cancer mutation-protein | Mutant Allele Frequency of background *TP53* mutations |  |  |
|  |  |  |  |  |  | Standard NGS MAF | Duplex Sequencing MAF | ddPCR MAF |
| con 2 | 70 | benign | - | c.733G>A | p.G245S | n.a. | 0.08% | 0.08% |
| con 7 | 79 | benign | - | c.817C>T | p.R273C | n.a. | 0.05% | 0.07% |

NGS: Next Generation Sequencing; MAF: Mutant Allele Frequency; ddPCR, digital droplet PCR; n.a: not assessed; <sup>†</sup> Spread to ovary *TP53* cDNA reference: NM\_000546.5, *TP53* protein reference: NP\_000537.3

Although *TP53* background mutations were common, their MAF was always below 1% (**Fig. 1E-F**), which could be used as a threshold to optimally identify patients with ovarian cancers from patients cancer-free. In this, albeit small, pilot study, a 1% threshold yielded a sensitivity of 70% and specificity of 100%, which outperformed other published tests for ovarian cancer detection (*8*). Furthermore, in the lavages where the tumor mutation was identified, its frequency was at least 10-fold above the highest background mutation in that individual.

### TP53 mutations in uterine lavage increase with age

To better understand the basis of *TP53* background mutations we examined the association of their abundance with age. When patients were ordered by ascending age (**Fig. 1E-F**), it appeared that older patients carried more mutations. However, the number of mutations found depends on total number of nucleotides sequenced (**Fig. S2**), which was variable across samples and tended to be higher in controls due to increased sequencing depths (**Table S2**). To compensate for this variation, for each sample we calculated the total *TP53* mutation frequency by dividing the number of mutations identified in uterine lavage (including exons and flanking intronic sequences) by the total number of Duplex bases sequenced. For patients with cancer, we excluded the tumor mutation from this calculation in order to fairly reflect only *TP53* background mutations. For patients with ovarian cancer, as well as cancer-free control patients, the *TP53* mutation frequency significantly increased with age (**Fig. 2**, p=0.0006 for ovarian cancer, p=0.001 for controls, Spearman’s correlation test). This trend was identical to prior observations of *TP53* background mutation frequency in peritoneal fluid and peripheral blood (*13*).

**Figure 2.**
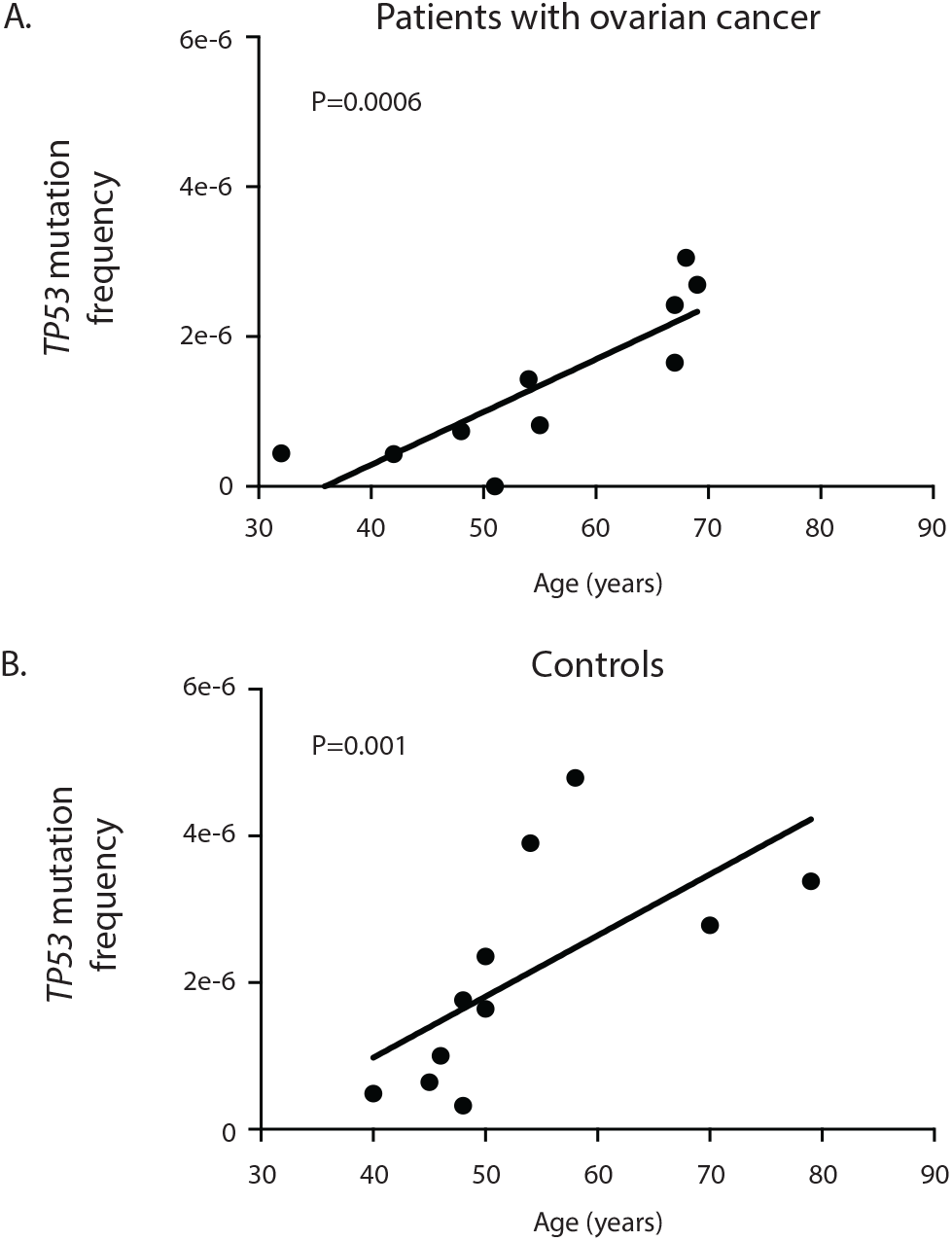
The frequency of TP53 mutations in uterine lavage increases with age. Frequency is calculated as the total number of unique TP53 mutations identified in each sample (including exons and flanking intronic regions) divided by the total number of Duplex nucleotides sequenced. (A) Uterine lavage samples from patients with ovarian cancer, n=10, r=0.89, p=0.0006 by Spearman’s correlation test. In these patients, the total count of TP53 mutations excludes the tumor mutation identified in the lavage in order to only represent TP53 background mutations. (B) Uterine lavage samples from control patients without cancer, n=11, r=0.83, p=0.001 by Spearman’s correlation test.

### TP53 mutations in uterine lavage are not random, but rather are positively selected

The *TP53* gene is a tumor suppressor, the genetic disruption of which facilitates cell proliferation in tumors, even when only one allele is mutated (*20*). An age-associated increase in ultra-low frequency *TP53* background mutations could result from random, age-related mutagenic processes or, alternatively, from mutagenesis coupled with clonal selection of pathogenic variants. To distinguish between these possibilities, we performed a detailed analysis of traits of selection among the 112 age-associated *TP53* background mutations collectively identified among all 21 patients (**Table S3**, **Fig. 3**).

**Figure 3.**
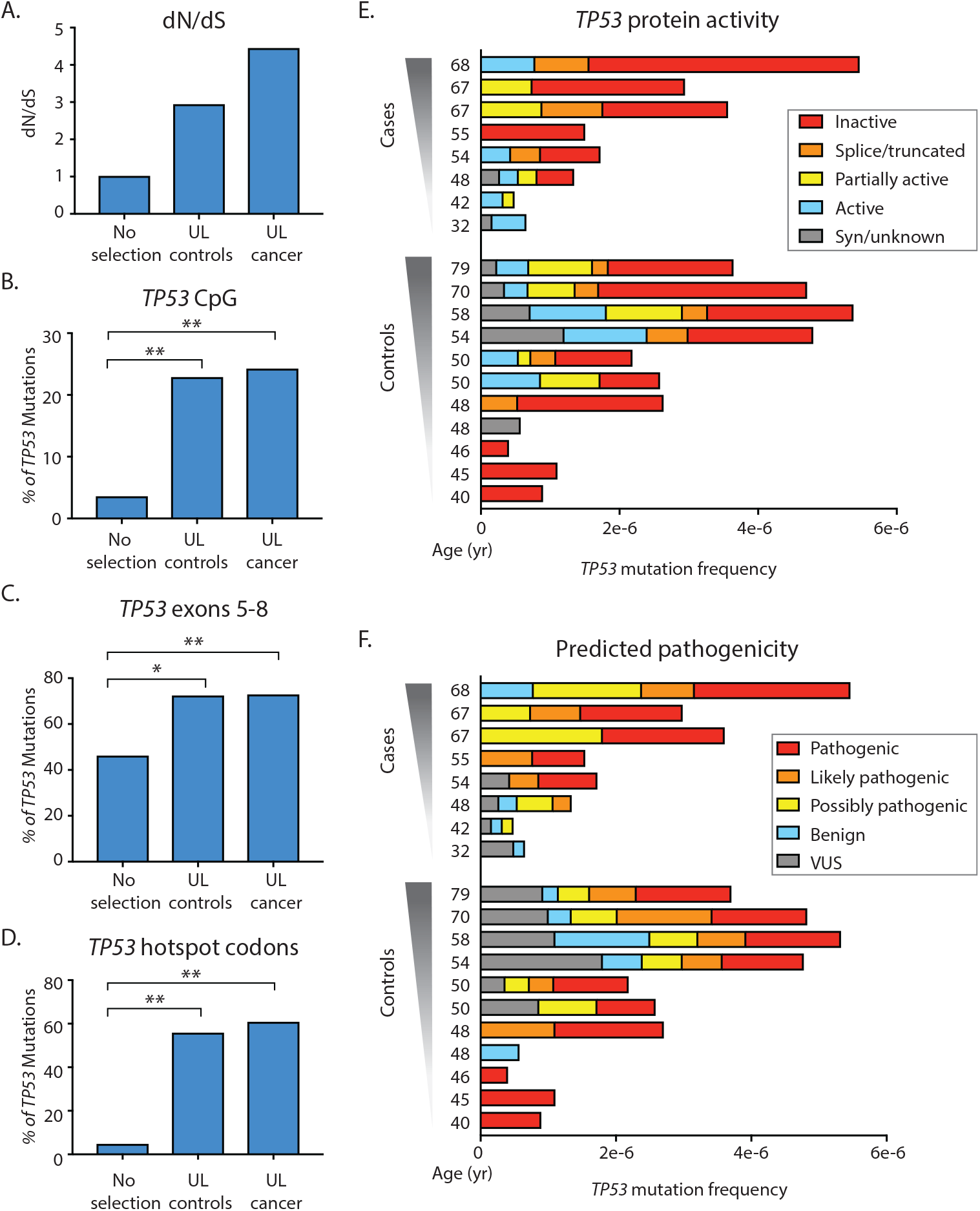
Evidence of positive selection in *TP53* background mutations from uterine lavages. (A) dN/dS. Values above 1 in uterine lavage from controls and cases correspond to an excess of non-synonymous mutations suggestive of positive selection. (B) Percent of *TP53* mutations localized in CpG dinucleotides. (C) Percent of *TP53* mutations localized in exons 5-8. (D) Percent of *TP53* mutations localized in hotspot codons. For C-D: *TP53* mutations identified in uterine lavage from controls and cancer significantly exceed expected values without selection. ^*^ p-value<0.01 ^**^ p-value<0.0001 by Fisher’s exact test, n=79 for uterine lavage controls and n=33 for uterine lavage cancer. (E) Protein activity and (F) predicted pathogenicity color-coded as 5 groups from Seshat data. Patients are sorted by ascending age. For each patient, *TP53* Mutation Frequency is calculated as the number of mutations in the coding region divided by the total number of Duplex nucleotides sequenced in that region. Two cancer patients with reduced sequencing depth and no identified *TP5*3 mutations are excluded from the analysis. Nearly all cases and controls carry mutations that have impact in protein activity and predicted pathogenicity. UL: Uterine lavage.

A metric of selection widely used in evolutionary biology, and recently incorporated into cancer genomics, is the dN/dS ratio (*21*). For a given coding region, this ratio compares the relative proportion of non-synonymous and synonymous mutations observed versus what would be expected based on random mutagenesis across all bases. A value of 1 indicates that the relative ratio observed is consistent with a random process and, in aggregate, coding changes in a region have neither a strong positive or negative _A._ impact on cell growth or survival. Values above and below 1, on the other hand, indicate positive and negative selection, respectively. The dN/dS ratio among background uterine lavage *TP53* mutations in cases and controls was 4.4 and 2.9, respectively, indicating enrichment for non-synonymous mutations that confer a growth or survival advantage (**Fig. 3A**). The excess of non-synonymous mutations was not driven by a subset of outlier samples, but rather was uniformly observed across nearly all lavage samples (**Fig. S3A**).

We next examined metrics of selection related to the genic location of mutations. Background *TP53* mutations were not randomly distributed along the gene but clustered in certain regions of biological significance. First, nearly a quarter of *TP53* lavage background mutations occurred in the context of methylation-sensitive CpG dinucleotides, which is remarkable given the fact that these dinucleotides comprise less than 5% of the coding region of *TP53* (**Fig. 3B**, p=5.8×10^-10^ for controls and p=1.7×10^-5^ for ovarian cancer mutations, by Fisher’s exact test). Mutations were also enriched in exons 5 to 8, which encode the DNA binding domain of the protein (**Fig. 3C**, p=3.3×10^-6^ for controls and p=0.002 for ovarian cancer mutations, by Fisher’s exact test). The most significant enrichment, however, was observed in *TP53* cancer-associated hotspot codons, which are the codons most recurrently observed mutated in cancer sequence databases. We considered the 9 most abundantly mutated codons in the UMD database (April 2017 version) (*20*). These codons encompass only 2.3% of the coding region of *TP53*, yet more than 25% of lavage background mutations clustered within these 27 base pairs (**Fig. 3D**, p=5.1×10^-17^ for controls and p=3.9×10^-9^ for ovarian cancer mutations, by Fisher’s exact test), and among these, all were non-synonymous. The biases seen for each characteristic were not driven by a subset of outlier samples, but were distributed homogeneously across samples in both groups (**Fig. S3B-D**).

To assess the impact of these mutations on *TP53* protein function, we took advantage of Seshat, a recently developed online tool for *TP53* analysis that provides comprehensive mutational information including prediction of impact on protein activity as well as pathogenicity (*22*). We queried all background *TP53* mutations identified in the 21 lavages and color-coded them according to 5 binned categories of protein activity and predicted pathogenicity. Nearly all samples carried at least one *TP53* mutation that inactivated the protein totally or partially (**Fig. 3E**) and/or was predicted to be pathogenic (**Fig. 3F**). The unambiguous signature across six distinct metrics of positive selection within the ultra-low frequency *TP53* mutations observed in all lavages, regardless of cancer status, indicate that these mutations expanded under strong selective pressure and are not the result of technical errors.

### TP53 mutations in uterine lavage resemble mutations in cancer

We next compared the features of selection of *TP53* mutations identified in lavages to *TP53* mutations from cancers. For this analysis, we used all the cancer mutations present in the UMD cancer database (April 2017, n=71,051). We determined the percentage of these mutations that reside at CpG sites, cancer hotspots, and exons 5-8, as well as the percentage of mutations that impact protein activity or are predicted to be pathogenic. For each trait, we compared the distribution of mutations in the theoretical absence of selection, in our 21 uterine lavages, and in the cancer database (**Fig. 4A**). Remarkably, for all traits, *TP53* background mutations from uterine lavages far more strongly resembled *TP53* mutations in the cancer database than the pattern expected by random chance. We also used a feature of Seshat that categorizes *TP53* mutations according to their frequency in the UMD database. Nearly all uterine lavage samples harbored *TP53* mutations listed as frequent or very frequent in the database (**Fig. 4B**).

**Figure 4.**
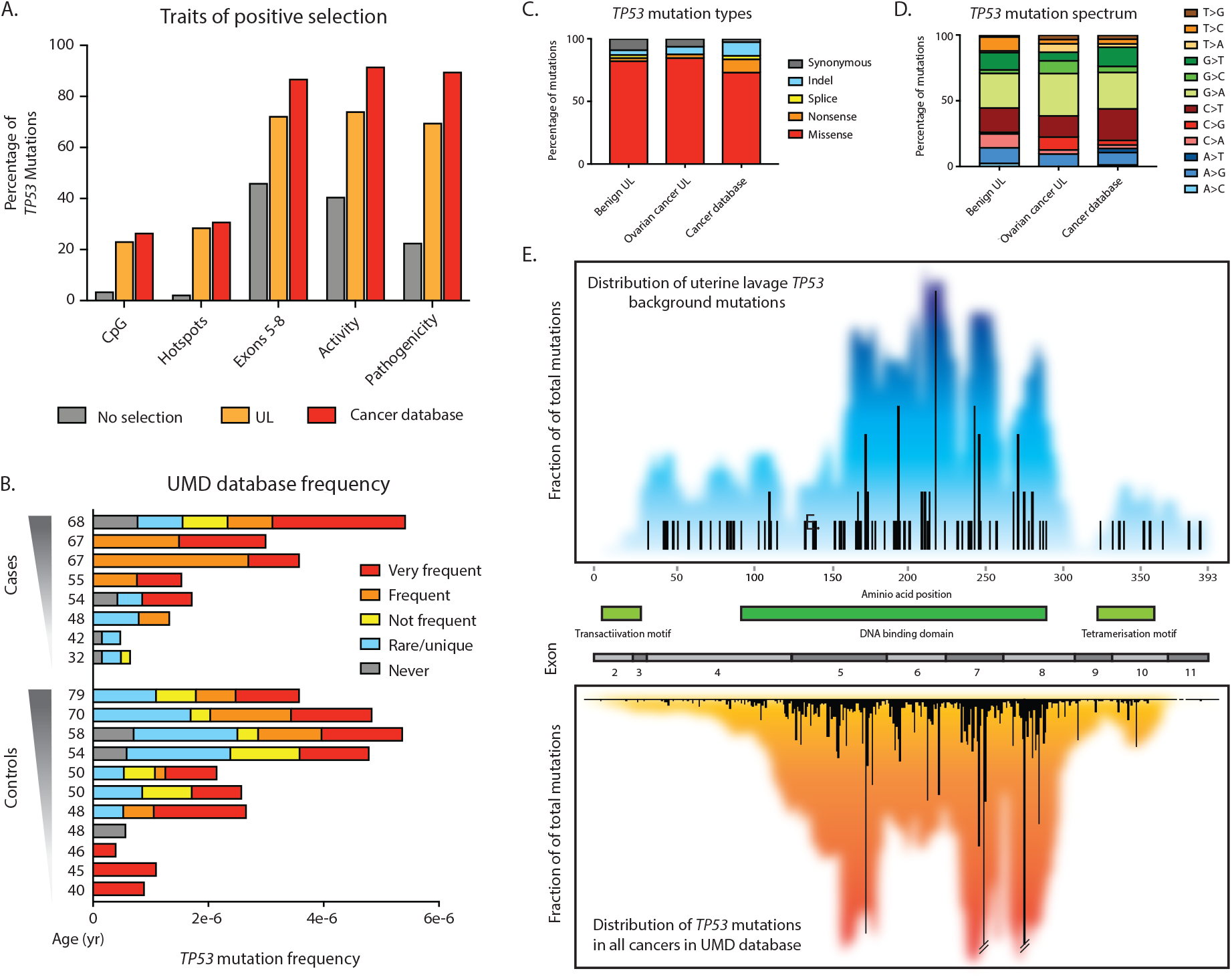
*TP53* mutations in uterine lavage are very similar to *TP53* mutations found in human cancers. (A) Traits of positive selection are compared between all possible mutations in the *TP53* coding region (No Selection, n=3,546), *TP53* background mutations found in uterine lavage (n=112), and *TP53* mutations reported in the UMD cancer database (n=71,051). (B) Uterine lavage mutations in cases and controls color-coded by their abundance in the UMD *TP53* database. For each sample, *TP53* Mutation Frequency is calculated as the number of mutations in the coding region divided by the total number of Duplex nucleotides sequenced in that region. Most samples harbor *TP53* mutations that are common in the database. (C) *TP53* mutation type and (D) *TP53* mutation spectrum are compared between mutations identified in uterine lavage from controls (n= 79), uterine lavage from cases (n=33) and the UMD database (n=71,051). (E) *TP53* mutation distribution map for uterine lavage (top panel) and UMD cancer database (bottom panel). Bars quantify the frequency of mutations at each codon. Colored background reflects a 20 BP sliding window average of mutation density around each position. Gene exons and protein domains are indicated in the middle section. UL: Uterine lavage.

To further characterize background *TP53* mutations in uterine lavage in comparison to those in cancers, we compared mutation type and spectrum distribution as well as location along the gene. Non-cancer derived mutations in uterine lavages from women with and without cancer were predominantly missense, similar to mutations in the database (**Fig. 4C**), and displayed a mutational spectrum enriched in G>A and C>T transitions, comparable to cancer mutations (**Fig. 4D**). Most strikingly, the distribution of low-frequency *TP53* background mutations from just 21 women along the length of the gene is a mirror image of the distribution of *TP53* mutations from more than 71,000 different tumors included in the database (**Fig. 4E**). Thus, somatic *TP53* mutations recovered from cells sloughed into the uterine cavity from normal healthy women are not random, but appear to emerge from an evolutionary process of mutation, selection and clonal expansion akin to what takes place in tumors, but within normal tissue.

### TP53 mutations are common in healthy tissues from middle age women

These striking results prompted us to consider what the tissue origin of the mutation-bearing cells in the uterine lavages might be. To address this question, we sequenced *TP53* from DNA obtained from pre-operative uterine lavage and peripheral blood as well as multiple gynecological tissues collected postoperatively following total hysterectomy/bilateral salpingo-oophorectomy for symptomatic fibroids (benign leiomyomas) from two middle age women (**Fig. 5A, Table S4**). DNA was extracted and processed for DS as previously, except that samples were sequenced to a higher average depth (**Table S5**).

**Figure 5.**
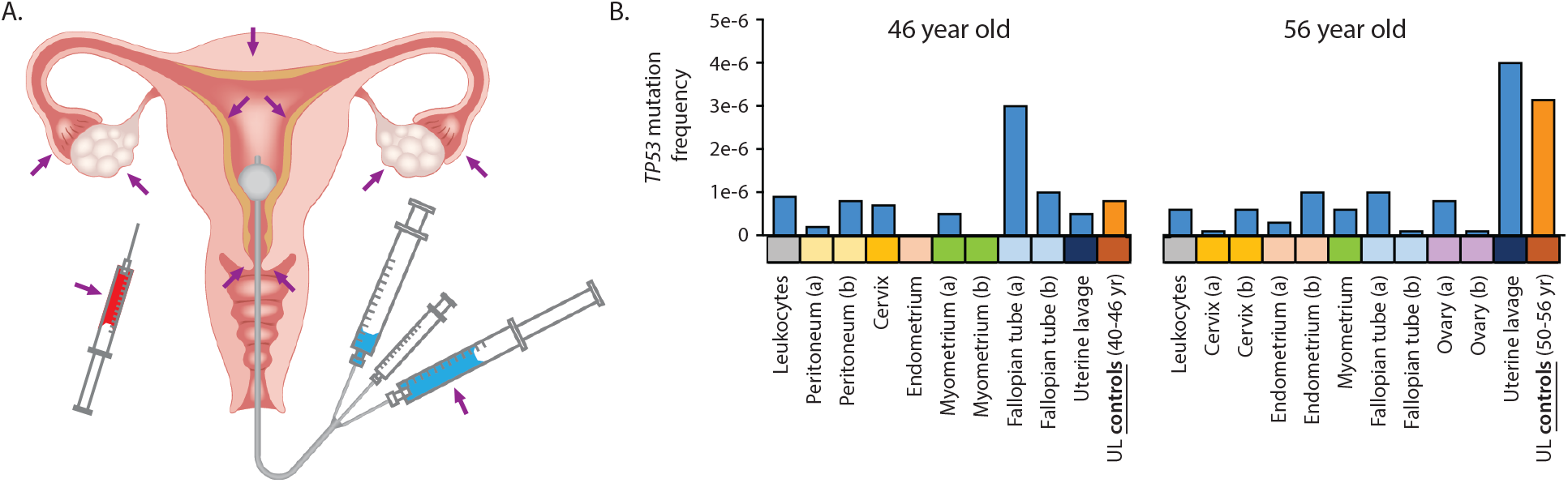
*TP53* mutations in normal tissues and uterine lavage from two middle age women. (A) Normal tissues collected including: leukocytes, peritoneum, cervix, endometrium, myometrium, fallopian tube, ovary, and uterine lavage. (B) *TP53* mutation frequency for each sample calculated as the number of *TP53* mutations in the coding region divided by the total number of Duplex nucleotides sequenced. a and b indicate two spatially separated samples from same tissue. Blue bars correspond to the samples in this study. Orange bars correspond to the mean values for uterine lavages from control women in the first study.

We identified *TP53* mutations in all samples from a 56 year old woman and in all but two samples from a 46 year old woman (**Fig. 5B, Table S6**). When we compared the mutation frequency across all samples, several interesting observations emerged. First, the uterine lavage from the 56 year old had a mutation frequency that appeared disproportionally high, both when compared to that of most other tissues and when compared with the lavage of the 46 year old. However, when compared to the mean values of uterine lavages from control women of similar ages (50-56 and 40-46 year old) from first part of the study, the frequencies by age were quite similar (**Fig. 5B**).

Moreover, the distribution of mutations according to each trait of positive selection (type, frequency in the cancer database, predicted activity and pathogenicity, exon clustering, CpG clustering, and enrichment for cancer hotspots) was comparable to the lavages previously analyzed (**Fig. S4**). Both lines of reasoning support the conclusion that the elevated frequency of mutations in the uterine lavage of the 56 year old is not artefactual and confirm the previously observed age effect.

For other tissues, however, we did not observe an obvious increase of *TP53* mutation frequency between 46 and 56 years of age. There was substantial variability in the mutation content of different tissues and of different biopsies of the same tissue, which reflects either a stochastic effect or the imprecision of macrodissection for obtaining exactly comparable tissue samples (for example, the depth of endometrium harvested or how distal the tubal fimbriae were cut). No single tissue stood out as obviously more mutation prone than another, nor could any tissue be identified as a dominant source of the mutations found in lavages.

### TP53 mutations increase in number and cancer-like features during normal human aging

With the hope of observing a stronger aging mutational signal, we looked to tissue samples from greater extremes of age. While the procurement of such material was challenging, we managed to obtain several gynecologic tissues at autopsy from a neonate who died from a congenital vascular malformation and from a 101 year old female who died of natural causes (**Table S4**). Together with the middle age samples, this unique specimen collection represents the full breadth of a century of the human lifespan.

Although the tissue types available were not fully identical across all four subjects, the pattern of *TP53* mutations, nevertheless, yielded unique insights. To help more intuitively visualize this multiparametric data, in **Fig. 6** we annotated all mutations found among the different tissues of the four subjects as color coded boxes for each feature of selection: red for “cancer-like”, blue for non-cancer-like. The number of columns of colored boxes per sample reflects the total number of mutations identified. When viewed in this format it is apparent that *TP53* mutations are not only more abundant with age but are also more “cancer-like”. Mutations found in older tissues are disproportionately observed in cancers and are predicted to inactivate the protein or be otherwise pathogenic. In contrast, mutations found in the newborn are rarely found in cancer, tend to preserve the protein activity, and are not predicted to be pathogenic.

**Figure 6.**
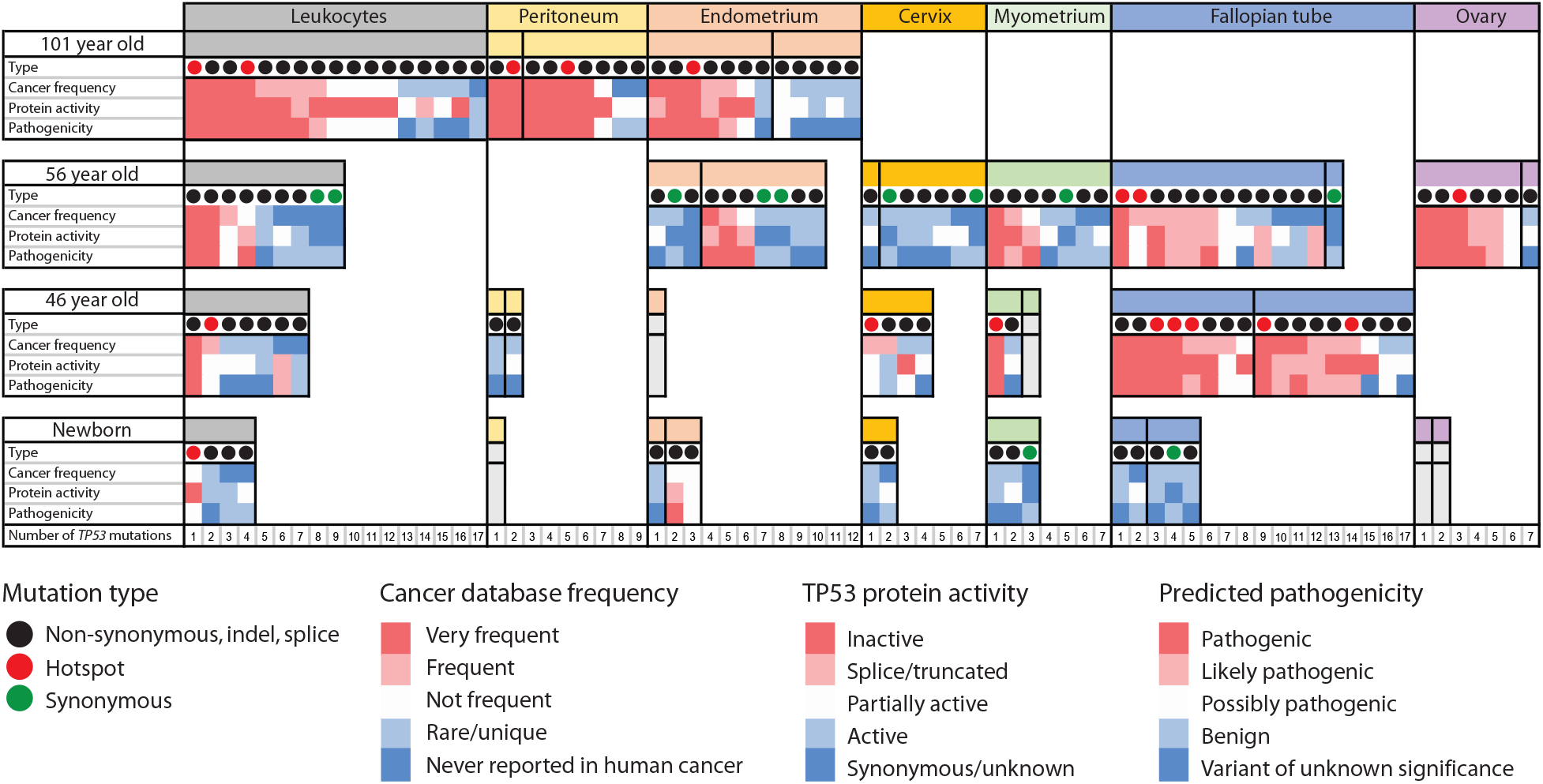
Characterization of *TP53* mutations in normal tissues over a century of the human lifespan. *TP53* mutations identified by DS in leukocytes and gynecological tissue are indicated as columns within each tissue. Each mutation is characterized by 4 parameters: type (synonymous, non-synonymous and hotspots); frequency in cancer; protein activity; and predicted pathogenicity. The last 3 parameters are color-coded with red indicating ‘cancer-like’ mutations and blue indicating benign mutations. Vertical black lines separate different biopsies from the same tissue. Within each biopsy, mutations are ordered left to right by decreasing cancer frequency. Biopsies that were sequenced but no mutations were identified are shown in grey.

Different tissues and different biopsies within the same tissue showed substantial variability in both the number of mutations and their cancer-like features. In aggregate, fallopian tube epithelium appeared to be a “hot” tissue with high number of mutations and a high percentage of cancer-like mutations. However, in the 56 year old, one fallopian tube sample harbored only a single synonymous mutation, consistent with the notion of “hot” and “cold” zones within a tissue. This was similarly seen in the two distinct endometrial biopsies of the centenarian.

In addition to a larger number of clones, with aging we would also predict an increase in the size of clones, as would be reflected by a higher MAF of each variant found. However, in this study, some samples were sequenced at a lower depth, which may lead to outlier (low event count) biases in the calculation of MAFs, thus precluding a fair comparison between samples (**Fig. S5**). Despite this caveat, two large clones were clearly seen within the peripheral blood leukocytes of the 101 year old (**Fig. S5** and **Table S6**, c.659A>G MAF: 1.2×10^-2^, and c.455C>T MAF: 4.5×10^-3^). Interestingly, the exact *TP53* mutation that defined each of these clones was also detected at lower frequencies in peritoneal and endometrial samples from the same subject, revealing an apparent contribution of leukocyte DNA to those tissue samples. (**Fig. S6**).

In fact, this cross-tissue mutation sharing was common in the 101 year old woman, suggesting that aged leukocytes might indeed harbor relatively large clones that recurrently contribute to the mutations found when sequencing other biopsies. Mutation sharing was less prevalent in middle age women. While very low frequency mutations are often hard to replicate due to the low precision of the measurement resulting purely from sampling statistics (not technical accuracy), it is important to keep in mind that certain mutations might also be recurrently identified simply because they are hotspots, and thus common origin cannot automatically be assumed. For example, the hotspot mutation c.659A>G was identified in the large blood clone of the 101 year old woman as well as in a myometrium sample and a fallopian tube biopsy of the 46 year woman (**Fig. S6**). The processing of these particular samples was done on different days, making a cross-contamination explanation improbable.

As already considered, an important limitation of this study was the different depth of sequencing achieved across samples, due to the inherent variability in library preparations as well as differences in DNA availability. Because numerically more mutations will be identified in samples with more sequencing (**Fig. S7A**), it is essential to compare samples based on their mutation frequency, which is a sequencing-normalized value calculated as the number of mutations divided by the number of total Duplex error-corrected nucleotides sequenced (**Fig. S7B**). *TP53* mutation frequency tended to be higher at older ages in the three tissue types shared by the neonate and the centenarian (leukocytes, peritoneum, and endometrium), although there was substantial variability across samples.

### TP53 mutations in newborn tissue are random, yet become positively selected over a lifetime

As further illustration of the increase of cancer-like mutations with aging, we divided all *TP53* mutations into two binary categories: “common in cancer” and “not common in cancer”, with the former being defined by those classified as “frequent” or “very frequent” in the UMD cancer database (**Table S7**). When plotted by age, the progressive enrichment for cancer-like mutations was easily apparent, especially in certain tissues such as fallopian tube and leukocytes (**Fig. 7A**).

**Figure 7.**
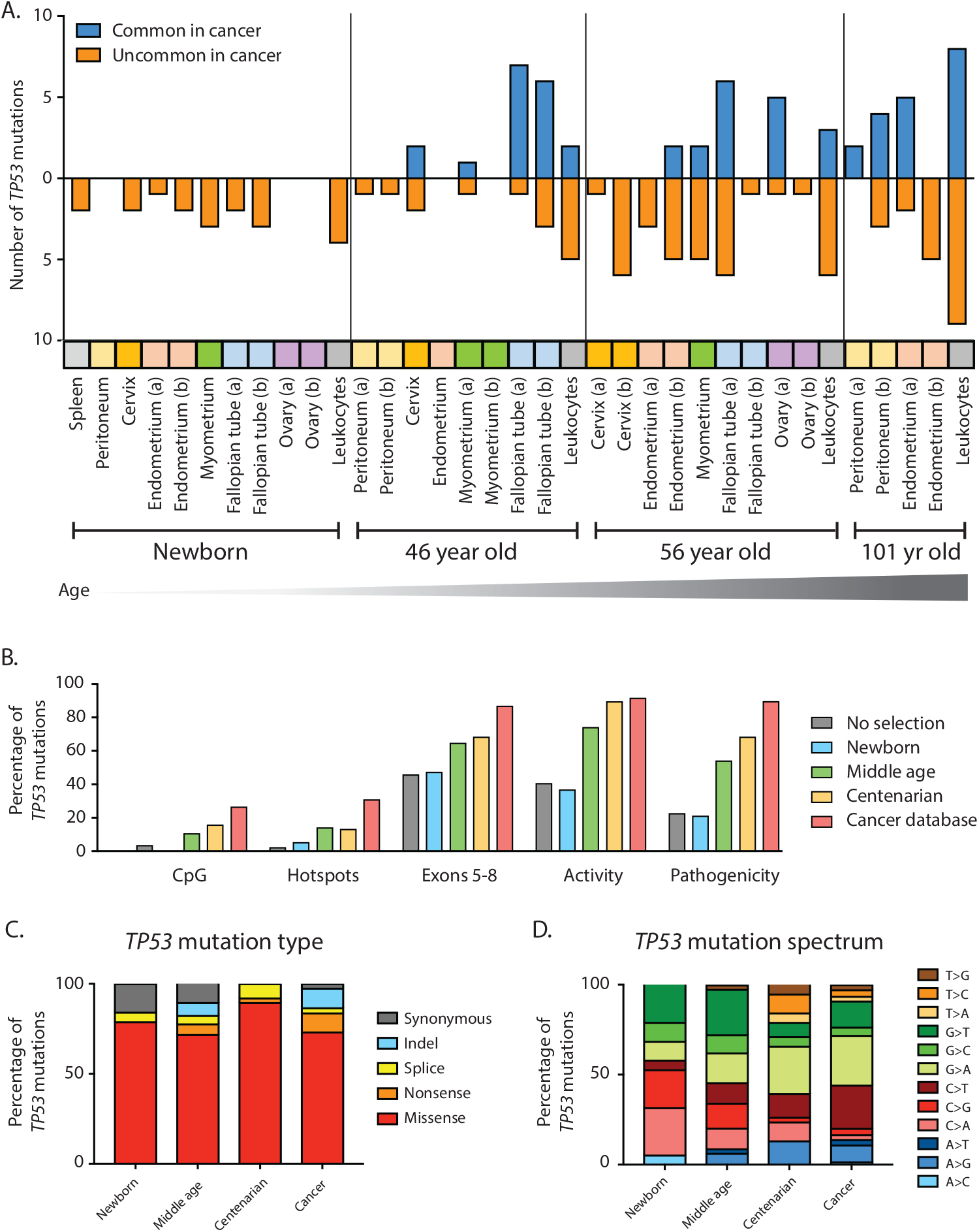
Cancer-associated *TP53* mutations are positively selected during normal aging. (A) Across a variety of human tissues, *TP53* mutations accumulate with age and are progressively enriched for mutations commonly found in cancers. Tissues are color-coded. ‘a’ and ‘b’ indicate two biopsies from the same tissue. (B) Traits of positive selection are compared between all possible mutations in the *TP53* coding region (n=3,546), *TP53* mutations found in newborn (n= 19), middle age (n=85), and centenarian (n= 38), and *TP53* mutations reported in the UMD cancer database (n=71,051). (C) Distribution of *TP53* mutation type and (D) mutation spectrum for newborn, middle age, and centenarian mutations (n=19, 85, and 38, respectively) compared to UMD cancer database (n=71,051).

We then examined the five traits of selection previously calculated for the uterine lavage study but for *TP53* mutations found in the newborn, middle age, and centenarian tissues (**Fig. 7B, Fig S8**). Remarkably, for mutations found in newborn, all five traits yielded values consistent with random processes (e.g. absence of selection), yet in middle age, and even more so in centenarian tissue, values reflected selection to an extent that neared that seen in mutations from tumors in the UMD database. Analysis of mutation type (**Fig. 7C**) revealed a decrease of synonymous mutations with age (in fact, no synonymous mutations were identified in centenarian tissue, **Fig. S8, Table S6**).

Lastly, regarding mutation spectrum, we noted an interesting preponderance of C and G mutations in newborn tissue, which progressively shifted towards an increased representation of A and T mutations in centenarian tissue, more similar to the pattern in cancers. The significance of this shift is unknown as it could represent both biases in the nucleotide composition of the gene at selectable hotspots, as well as differential age-associated mutagenic processes (*23*) which disproportionately contribute to the clonal mutation burden of tumors because tumors mostly arise in the elderly.

### TP53 mutations in cfDNA and peritoneal fluid follow the same patterns as solid tissue

To explore the abundance of *TP53* mutations to liquid biopsies of clinical interest, we analyzed plasma-derived cell-free DNA (cfDNA) and peritoneal fluid from the 46 year old woman. *TP53* mutations were identified in both, with cancer-like features similar to what was observed for solid tissue biopsies and leukocytes (**Fig. S9**). None of the mutations identified in cfDNA or peritoneal fluid overlapped with mutations identified in leukocytes, uterine lavage or any of the solid tissues analyzed (**Fig. S6).** Peritoneal fluid is routinely collected for disease staging during gynecological surgery and we previously demonstrated that it carries *TP53* background mutations with cancer-like features (*13*). cfDNA had not been analyzed previously by DS. The fact that one of the identified mutations is pathogenic and commonly found in cancers (**Fig. S9, Table S6**) raises important concern over specificity in cancer-screening studies based purely on mutation detection in plasma.

## DISCUSSION

We have demonstrated that uterine lavage coupled with DS offers a promising solution for ovarian cancer detection based on a minimally invasive sampling approach that can practically be integrated into routine gynecologic primary care (*24*). The technique improves upon past mutation-based screening efforts through use of (1) a collection method that recovers cancer cells very close to the anatomical site of the tumor and (2) an ultra-accurate DNA sequencing technology that can resolve exceptionally low frequency mutations. In this small study, we were able to achieve remarkable sensitivity and specificity using a mutation allele frequency threshold for differentiating cancer cases from non-cancer controls. This was possible without the allele-specific PCR technique required in our past lavage study (*9*) which, from the perspective of a screening test, impractically necessitated prior knowledge of the specific tumor mutation being sought. While validation through substantially larger prospective trials will be critical to support widespread clinical use, we have established the technical foundation for such studies, which are now enrolling in both Europe and the US (LUDOC II - ClinicalTrials.gov Identifier: NCT02518256, LUSTIC - ClinicalTrials.gov Identifier: NCT02039388).

However, the most profound finding of this work is not the biomarker performance of the technique itself, but the incidentally found mutational patterns that reflect a somatic evolutionary process that appears operative throughout much of human life in normal tissues. Specifically, we identified widespread low frequency *TP53* mutations that were heavily enriched for pathogenic variants. This enrichment reflects a process of natural selection that favors the survival and proliferation of cells with mutations that are identical to those observed in cancer, but as part of routine aging. The unambiguous selection signature is supported by multiple orthogonal metrics and cannot be explained by technical errors; both the biological and diagnostic implications are substantial.

One of the main reasons that cancer biomarkers fail to reach the clinic is their inability to achieve the extremely high specificity required for screening (*25*). This is critical for cancers with low incidence and that require an invasive procedure to follow up positive screening tests, such is the case of ovarian cancer (*3*). Harms due to false positives and a lack of proven reduction in mortality are the reasons for the recent recommendation against the use of CA-125 and transvaginal sonography for screening asymptomatic women (*4*). In recent years, DNA mutation-based cancer screening from plasma or other body fluids has emerged as a promising method to detect cancer based on the supposition that cancer-associated mutations found in liquid biopsies are a specific indication of cancer somewhere in the body (*26*). Here we demonstrate that with a sufficiently low technical background, biologically real cancer-associated mutations can be found in every tissue tested: cancer-associated mutations are, in fact, far from cancer-specific.

The detection of cancer-associated mutations in normal tissues is not entirely new (*14*). In 2014, a series of major publications reported that mutations commonly found in acute myeloid leukemia occur as minority subclones in the blood of ~10% of healthy elderly individuals–a phenomena dubbed Clonal Hematopoesis of Indeterminate Potential (CHIP) (*27-29*). A year later Martincorena and colleagues observed hundreds of tiny clones carrying cancer-associated driver mutations on sun-exposed eyelids (*30*), a finding recently replicated in normal aged esophagus (*31*). Cancer-associated mutations have been similarly reported in abnormal, but non-cancerous tissues including endometriosis (*32, 33*) and benign dermal nevi (*34*). The use of laser capture microdissection in recent studies has revealed that as many as 1% of normal colorectal crypts of middle-age individuals (*35*) and >50% of normal endometrial glands of middle-age women (*33*) carry mutations in cancer driver genes.

The relationship between oncogenic mutations and cancer has been known for decades, yet the delay in appreciating their presence outside of cancer can be largely attributed to technical limitations. The advent of NGS enabled surveying wide swaths of the genome and detection of mutations present clonally or as modest size subclones. In the above studies, standard whole exome or multi-gene NGS were able to identify driver mutations because of unique scenarios where clones were either relatively large (CHIP in a subset of very elderly) (*27, 28*) or spatially coherent and comprising a sizeable percentage of cells when very small biopsies were taken (*30, 33, 35*). With higher accuracy NGS techniques able to resolve lower frequency subclones, later studies have found CHIP mutations at lower levels in most middle-aged adults (*36, 37*). Using ultra-high accuracy DS, we found extremely low frequency cancer-associated *TP53* mutations in both blood and peritoneal fluid of healthy women undergoing abdominal surgery and, in both sample types, the abundance of mutations increased with age (*13*). A subsequent study that employed uterine lavage for endometrial cancer detection found pathogenic mutations in cancer driver genes in lavages of cases as well as controls (*38*).

The present study adds a new layer of knowledge by identifying extremely low frequency *TP53* mutations in uterine lavages and multiple normal tissues. DS has an error rate below one-in-ten-million, so even mutations that are seen in a single nucleotide among hundreds of millions of other sequenced nucleotides could be confidently identified. In aggregate, we sequenced 319,576,913 unique *TP53* coding nucleotides and identified 292 non-cancer derived “biological background” mutations. The average background mutation frequency of 9×10^-7^ in this study is at least ten thousand times below the background error threshold of standard NGS and dozens of times below that of other error correction methods (*10*) (**Fig. S1**).

We examined 10 different tissue/sample types from a unique cohort of individuals spanning more than a century of the human lifespan and assessed the pattern of mutations found using multiple different metrics of selection. A significant and novel finding was that, not only do *TP53* mutations increase in abundance with age, but the relative representation of random mutations versus cancer-associated mutations transitioned from almost entirely the former to almost entirely the latter from birth to end-of-life. Moreover, the extent of mutation frequency increase varied considerably by sample type and tissue. Uterine lavage samples carried nearly an order of magnitude more mutations than any other tissues examined by a woman’s mid 50’s. The basis of the rapid increase from women in their mid 40’s to those in their mid 50’s could plausibly relate to the onset of menopause. The majority of cells collected in uterine lavage are of endometrial origin and the cessation of cyclic sloughing at menopause might eliminate a physiologic means of purging mutated cells. Of note, none of the 10 *TP53* coding mutations identified in the endometrium of the 56 year old woman was detected in the uterine lavage, which contained as many as 25 mutations. This could be explained by the fact that the endometrial biopsies analyzed contained not only superficial surface epithelium, which are the cells collected by the lavage, but deep tissue as well, which might have a lower mutational load.

The biology seen here is likely only the tip of the iceberg and much further work remains to be done. We only examined tissue sets from four individuals at a very coarse spacing along the aging continuum. We focused on only gynecologic tissues and did not have perfectly matching samples for each subject, given how challenging these unique specimens were to obtain at the extremes of age. Furthermore, we only sequenced the coding region of a single gene, albeit the one most commonly mutated in cancer.

The implications of our findings are important as related to the physiology of aging, but also as a cautionary message for mutation-based cancer biomarkers of all varieties. At the same time as we have shown that high sensitive NGS methods are essential for maximally sensitive mutation-based cancer diagnosis, we have also illustrated a substantial specificity challenge related to biology, not technology, the extent of which has been under-appreciated. This is not limited to one or a few tissues, rather, it seems to be ubiquitous among the epithelial, mesenchymal, and hematopoietic cells lineages we investigated. Moreover, the same selection patters of mutations were found in minimally invasively collected body fluids, including both uterine lavage and cfDNA. This suggests that ongoing large scale efforts to develop universal “liquid biopsy” cancer screening tests via deep sequencing of cfDNA from plasma need to be approached with great caution (*26, 39*).

Despite the extent of biological background mutations, as a non-invasive cancer test our approach worked remarkably well. We identified 80% of tumor mutations and 70% of those were above the 1% MAF threshold we used to distinguish cases from controls. The only tumor mutation missed below this threshold corresponded to a 42 year old woman, one of the youngest in the study. Younger women tended to have fewer background mutations and mutations with lower MAF, which suggest that, moving forward, sensitivity could be increased by using age-adjusted thresholds. In addition, specificity could be improved by: uniform lavage collection at the luteal phase in premenopausal women, sequencing of peripheral blood to identify and exclude CHIP clones that might be present in lavage and longitudinal assessment of mutations to identify MAF increases over time. An important consideration is that we have demonstrated detection of intermediate and late stage cancers, but the most critical target for screening are early stage cancers because they are most curable. In that regard, monitoring of high risk populations, such as *BRCA1* and *BRCA2* carriers, may be the highest impact near-term clinical implementation.

The sensitivity improvements lent by new sequencing technologies are forcing a far more nuanced genetic definition of what distinguishes a cancer cell from simply an old cell. Our results show that CHIP clones are merely one relatively easy-to-detect manifestation of a far broader phenomena that appears to extend to most, if not all, tissues in the body. From a biomarker perspective, the fact that those who are at greatest risk of cancer and for whom cancer screening holds the most benefit (older adults) are also the population with the most cancer-like age-associated background mutations, is particularly inconvenient. Ongoing improvements will be needed to find ways to maximize specificity through careful MAF threshold calibration and combination with orthogonal biomarkers. Further investigation into the significance of biological background mutations from the perspective of human aging and biology is similarly warranted. For example, does the frequency of mutations and extent of pathologic shift provide a chronologically-independent empiric measure of age? Could it be used to integrate a lifetime load of intrinsic and extrinsic mutagenic exposures and predict future cancer risk?

While the notion that our somatic genomes are steadily evolving towards neoplasia with each passing decade might viewed as disheartening, an alternative perspective is that, in spite of this, most people do *not* develop overt cancer in their lifetime. This serves as a reminder of just how much remains unknown about the body’s many complex mechanisms of tumor suppression–a toolkit that we can perhaps augment with future technologies for cancer prevention.

## MATERIAL AND METHODS

### Experimental Design

We performed two complementary studies. The objective of the uterine lavage study was to determine the ability of DS to detect ovarian cancer through deep sequencing of *TP53* mutations in uterine lavage. This study included 10 patients with high grade serous ovarian cancer (cases) and 11 with benign gynecological masses (controls). The objective of the normal tissue study was to characterize somatic *TP53* mutations that accumulate during aging. This study included tissue from two newborn subjects (one newborn male only provided blood), two middle age women (ages 46 and 56 years old) and one centenarian woman (101 years old). Clinico-pathological information for all subjects is listed in **Table S1**.

### Samples

In the first study, we analyzed uterine lavages collected by a trans-cervical catheter (**Table S1**). Lavages were collected immediately pre-operatively as previously described (*9*). Lavage samples were centrifuged at 300x g for 10 minutes at room temperature and DNA was isolated from the cell pellet (QIAamp MinElute Kit, Qiagen, Hilden, Germany). Patients were recruited in three institutions: Medical University of Vienna (Austria), Charles University Pilsen (Czech Republic) and University Hospitals Leuven (Belgium). Sample procurement was performed in accordance with the institutional review boards of the Medical University of Vienna (EK#1148/2011 and EK#1766/2013), the Catholic University Leuven (B322201214864/S54406) and the Medical Faculty Hospital Pilsen (No 502/2013).

In the second study, multiple gynecological tissues were collected per **Table S4**. Not all sample types were available for all subjects. Newborn and centenarian tissue was collected at autopsy while tissue from middle age women was collected following hysterectomy. Peripheral blood was unavailable from the female newborn and to compensate we sequenced spleen from this subject as well as peripheral blood sample from a neonatal male (7 weeks old). The two newborn autopsies were performed at Seattle Childrens’ Hospital and tissues were collected under research IRB #52304. For the two middle age women, uterine lavage was collected with the same procedure as in the first study. In addition, for the 46 year old woman, cfDNA was collected preoperatively and peritoneal lavage collected intraoperatively. Both operations were performed at the Medical University of Vienna and samples were collected with informed consent and according to approved IRB EK# 1152/2014. For the centenarian case, tissue was obtained via rapid autopsy from Tissue for Research Inc. and processed at the University of Washington under IRB waiver 2016-52304. All samples were collected using sterile new instruments between biopsies and frozen over liquid nitrogen immediately after collection and stored at −80°C until DNA extraction.

### Digital Droplet Polymerase Chain Reaction

Lavage DNA from 5 ovarian cancer cases and 2 controls was analyzed by ddPCR (**Table 1**). In ovarian cancer lavage, ddPCR amplified the tumor mutation whereas in benign lavage, the assay targeted two mutations previously identified by DS at frequencies below 0.1%. ddPCR was performed with the QX100 Droplet Digital PCR system (Bio-Rad Laboratories, Hercules, CA) using custom TaqMan SNP Genotyping Assays (Life Technologies, Carlsbard, CA) designed using Primer Express 3.0 software (ThermoFisher). 10-20ng of DNA were used in each reaction and samples were analyzed at least in duplicates. A positive control and a wild-type control were included in every run.

### Duplex Sequencing

Duplex Sequencing was performed as previously described with minor modifications (*12*). Briefly, DNA was sonicated, end-repaired, A-tailed, and ligated with DS adapters using the KAPA HyperPrep library kit (Roche Sequencing, Pleasanton, CA). After initial amplification, 120 bp biotinylated oligonucleotide probes (Integrated DNA Technologies, Coralville, Iowa) were used to capture the coding region of *TP53*. Two successive rounds of captures were performed to ensure sufficient target enrichment, as previously described (*40*). Indexed libraries were pooled and sequenced on an Illumina HiSeq2500 or NextSeq 550. Sequencing reads were aligned to hg19 then reads sharing a common molecular tag in both distinct strand orientations were grouped and assembled into an error-corrected Duplex Consensus Sequence as previously described (*12*).

The total number of Duplex nucleotides sequenced for each uterine lavage and tissue sample is listed in **Tables S2** and **S5**. In aggregate, we sequenced 587,169,708 unique nucleotides, 319,576,913 of which corresponded to coding nucleotides. We targeted a median Duplex depth of ~1000x. Three tissue biopsies were excluded because of insufficient depth. Because Duplex reads correspond to original DNA molecules, Duplex depth indicates the total number of haploid genomes sequenced. For each sample, *TP53* mutation frequency was calculated as the number of identified mutations divided by the total number of Duplex nucleotides sequenced. For each individual mutation, mutant allele frequency (MAF) was calculated as number of mutated Duplex bases divided by the total Duplex depth at a given nucleotide position. Mutations identified as SNPs in the 1000 genome database with were excluded from mutation analysis. All mutations were manually reviewed with the Integrative Genome Viewer (IGV).

### Characterization of TP53 mutations using Seshat and the UMD TP53 database

The final list of mutations from all samples in the study (uterine lavage study n=166, normal tissue study n=264) was converted into a Variant Call Format (VCF) file and submitted to Seshat (https://p53.fr/TP53-database/seshat), a web service that performs *TP53* mutation annotation using data derived from the UMD *TP53* database (*22*). This database is the most updated and comprehensive repository of *TP53* variants. From the Seshat output (included as **Database 1** for uterine lavage mutations and **Database 2** for normal tissue mutations), the following variables were extracted: cDNA variant, HG19 Variant, Variant Classification, Frequency, Activity, Pathogenicity, Exon, Codon, CpG, Mutational event and Variant Comment. These variables were used to annotate the DS pipeline-generated mutational calls in **Table S3** (uterine lavage) and **Table S6** (normal tissue). The human genomic reference hg19 (GRCH37) was used for data reporting. Mutations occurring in the coding region and adjacent splice sites were selected for mutational analysis (uterine lavage n=112, normal tissue n=180).

Mutations were annotated based on type (missense, nonsense, splice, synonymous), mutation spectrum (each of the 12 possible nucleotide substitutions), localization to CpG dinucleotides, localization in exons 5 to 8 (encoding the protein’s DNA binding domain), localization to mutational hotspot (9 most common mutated codons in the UMD *TP53* database: 175, 179, 213, 220, 245, 248, 249, 273, 282), frequency of the mutation in the cancer database, functional activity, and predicted pathogenicity. Functional activity was assessed by a transcriptional activity chart assay for 3,000 variants performed by Kato et al. (*41*). Pathogenicity was based on multiple predictive algorithms included from dbNSFP (*42*) as well as functional activity (*22*). For the last 3 variables (frequency in cancer database, activity and pathogenicity), mutations were aggregated into binned groups (**Table S7**).

### TP53 cancer database mutational analysis

From the UMD *TP53* database (April 2017 version), we selected the set of 71,051 mutations reported within human tumors of all types (mutations from cell lines, normal and premalignant tissue were excluded). Then we determined the distribution of mutations in the following categories: CpG, hotspots, exons 5-8, activity, pathogenicity, mutation type and mutation spectrum. These values were used as a comparator for *TP53* mutations identified in uterine lavage and normal tissues.

### TP53 mutations without selection

To assess the distribution of *TP53* mutations in the absence of selection, we generated a list of all possible mutations in the gene coding region (n=3,546) *in silico*. Then we submitted this list to Seshat to determine the distribution of mutations in the same categories as above. The values obtained represent the distribution of all possible *TP53* mutations in the absence of selection and were used as a comparator for *TP53* mutations identified in uterine lavage and normal tissue.

### dN/dS calculation

The dN/dS ratio measures the ratio of non-synonymous vs synonymous mutations taking into consideration observed and expected values for a given genomic region. It was calculated as (n/s)/(N/S), where n is the total number of observed nonsynonymous mutations, s is the total number of observed synonymous mutations, N is the total number of possible nonsynonymous mutations, and S is the total number of possible synonymous mutations. For *TP53*, N=2,715 and S=831 based on the 3,546 possible mutations in its coding region. A dN/dS ratio equal to 1 indicates neutral selection, <1 indicates negative selection, and >1 indicates positive selection.

### Statistical analyses

Correlations were tested with Spearman’s rank test due to high variability in the outcomes (nonnormality). The comparison of *TP53* mutational traits in uterine lavage of controls and cancers *vs* non-selected mutations was performed with Fisher’s exact tests. All tests were two-sided at an alpha level (type 1 error rate) of 0.05. Statistical analyses were performed with SPSS and R.

## Acknowledgments

We thank members of the Loeb, Risques, Kennedy, and Swisher labs at the University of Washington for helpful discussions as well as participating members of the LUSTIC and LUDOC clinical trials. Most importantly we deeply thank the patients and families who volunteered to provide clinical samples, without which this research would not have been possible.

## Author contributions

JJS, RZ, PS, EM and RAR designed the study; EM, EL, RH and AV procured the samples; JJS, KLS, EM, and LNW processed the samples; JJS, CCV, DN, KB, TS and RAR contributed to data analysis and visualization; ME and RAR performed statistical analyses; LAL, RZ, and PS contributed expertise and invaluable critical discussion; JJS and RAR wrote the article.

## Data and materials availability

Sequencing data that supports the findings of this study have been deposited in the Sequence Read Archive (BioProject ID: PRJNA503496). Software for DS data analysis is available at https://github.com/risqueslab.

## SUPPLEMENTARY MATERIALS

**Figure S1**. Comparison of mutation detection limit by sequencing accuracy for different NGS methods

**Figure S2**. Association between number of independent *TP53* mutations detected and total number of Duplex nucleotides sequenced

**Figure S3**. *TP53* mutation frequency and characteristics by age for individual patient lavages in case-control study

**Figure S4**. *TP53* mutation frequency and characteristics by age including uterine lavages from the two middle age women in the normal tissue study

**Figure S5**. Mutant allele frequency as a function of Duplex sequencing depth

**Figure S6**. Analysis of mutations shared across multiple tissue samples within the same individual

**Figure S7**. *TP53* mutation frequency by tissue type

**Figure S8**. *TP53* mutation frequency and characteristics by age for individual tissue samples

**Figure S9**. *TP53* mutation characteristics within non-invasively collected body fluids

**Table S1**. Clinico-pathological characteristics of patients

**Table S2**. Uterine lavage Duplex Sequencing coverage

**Table S3**. *TP53* mutations detected by Duplex Sequencing in uterine lavage

**Table S4**. Clinico-pathological characteristics of individuals that provided normal tissue

**Table S5**. Normal tissue Duplex Sequencing coverage

**Table S6**. *TP53* mutations detected by Duplex Sequencing in normal tissue

**Table S7**. Postprocessing of Seshat analytical variables into categorical variables

**Database S1**. Seshat’s long form analysis of *TP53* mutations identified by DS in uterine lavage

**Database S2**. Seshat’s long form analysis of *TP53* mutations identified by DS in normal tissue

